# The effect of education on adult mortality, health, and income: triangulating across genetic and policy reforms

**DOI:** 10.1101/250068

**Authors:** Neil M Davies, Matt Dickson, George Davey Smith, Frank Windmeijer, Gerard J van den Berg

**Author notes:** **Classification:** Social Sciences (Economic Sciences) and Biological Sciences (Genetics). **Conflicts of interest:** We report no conflicts of interest.

## Abstract

On average, educated people are healthier, wealthier and have higher life expectancy than those with less education. Numerous studies have attempted to determine whether these differences are caused by education, or are merely correlated with it and are ultimately caused by another factor. Previous studies have used a range of natural experiments to provide causal evidence. Here we exploit two natural experiments, perturbation of germline genetic variation associated with education which occurs at conception, known as Mendelian randomization, and a policy reform, the raising of the school leaving age in the UK in 1972. Previous studies have suggested that the differences in outcomes associated with education may be due to confounding. However, the two independent sources of variation we exploit largely imply consistent causal effects of education on outcomes much later in life.

## 2 Introduction

Educational decisions, such as choosing to remain in school, made comparatively early in life associate with substantial differences in outcomes across the life course.(*1*–*8*) Unfortunately for researchers interested in the causal effects of education, these choices do not occur at random. For example, on average people who chose to remain in school for longer are more likely to have educated parents. Thus it is challenging to determine if education causes differences in outcomes later in life, or if other, potentially unknown, factors drive these associations. As a result, approaches such as multivariable adjustment are likely to suffer from residual confounding.(*9*) In contrast, instrumental variable analysis can potentially estimate the causal effects of education in the presence of unmeasured confounding of the education-outcome association. Three assumptions define instrumental variables: 1) they must associate with the risk factor of interest (the “relevance/informativeness criterion”); 2) they have no common cause with the outcome (“the independence assumption”); and 3) they have no effect on the outcome except via the risk factor of interest (the “exclusion restriction”).(*10*) Natural experiments, such as legal changes to school leaving ages are potential instrumental variables for educational attainment. These changes forced people to remain in school for longer, and, because parents could not have anticipated them, are unlikely to be associated with factors that confound the association of education and other outcomes. The size of the effect of education can be estimated using instrumental variable estimators.(*1*)

Another potential instrumental variable for education are genetic variants that are known to associate with educational attainment.(*6, 11, 12*) The use of genetic variants as instrumental variables is known as Mendelian randomization. This approach exploits the natural experiment that occurs at conception – when each child inherits half of each of their parents’ genomes. This process means that at each locus there is a 50% chance of inheriting one or other of their parents’ alleles. The first instrumental variable assumption is likely to hold because large genome-wide association studies (GWAS) have discovered genetic variants that robustly associate with education. Because of the segregation of alleles at conception, these genetic variants are also independent of many confounders. While many phenotypes are far more associated with each other than would be expected by chance, genetic variants known to associate with one trait, tend to be independent of other potential risk factors.(*13*) Furthermore, each person’s genome is set at conception cannot be affected by their later educational choices or other outcomes.

While legal changes to school leaving ages have been widely used as an instrumental variable for education, genetic variants known to associate with education have received less attention. The instrumental variable assumptions are plausible for phenotypes whose biological pathways are relatively well understood (e.g. variants in the CRP gene for CRP levels(*14*) or variants in ALDH2 for alcohol(*15*)), they may be less plausible for phenotypes where the mediating pathways are less well understood such as education. For example, genetic variants that affect parents’ education may have direct effects on the offspring (so-called “dynastic effects”); parents assortatively mating on education(*16*); or that more educated parents have different ancestry from those with less education. These potential sources of bias are illustrated in **Supplementary Figure 1.** While Mendelian randomization using samples of unrelated individuals may be a credible identification strategy for biologically proximal phenotypes such as CRP or alcohol consumption, it may be less plausible for biologically distal phenotypes such as education. Recent studies have used genetic variants known to associate with education to estimate the effects of education on coronary heart disease and dementia.(*6, 12*) However, unlike hypotheses that relate to biological traits, such as lipids, there are no randomized trials that can provide gold-standard evidence of the causal effects of education.

Two key questions in the scientific and policy literature, are whether the effects of education across individuals or at different points in the life course are heterogeneous.(*17*–*20*) For example, does an additional year of schooling at age 16 have the same effect on everyone? Does an additional year of schooling at age 16 have the same effect as an additional year of schooling at age 20? Many of the previously investigated policy reforms affect a subset of individuals at a specific age (e.g. the effect of an additional year of education for low ability students at age 16). Policy makers may be interested in the effects of education on average across the whole population, or of the effects of obtaining a specific length of schooling (e.g. staying in school to age 18 versus 16). However, genetic variants affect educational choices across the entire lifespan. They identify the average effect of an additional year of school across the entire cohort.

Here we compare two potential instrumental variables, a policy reform and Mendelian randomization within the same sample. We have previously reported the effects of educational attainment using the raising of the mandatory minimum school leaving age using data from the UK Biobank.(*21*) We assess the plausibility of the Mendelian randomization assumptions for estimating the effects of educational attainment. We estimate the long-term effects of education using both genetic variants and the raising of the school leaving age.

## 3 Results

### 3.1 Descriptive statistics

The UK Biobank invited 9.2 million people aged between 40-69 to attend 23 centres across Great Britain.(*22*) Of those invited 503,325 (5.47%) were recruited in 2006-2010 to the study. Of these 315,436 met the inclusion criteria for this study. See the supplementary materials for a flowchart of the inclusions and exclusion of participants (**Supplementary Figure 2**). The average age when attending the assessment centre was 56.9, and 53.8% were female. On average, UK Biobank participants were more educated than the British population, 41.0%, 64.0%, and 82.1% had a degree or equivalent, had post-16 education, and any academic qualifications respectively. Whereas the UK census found that 27.9%, 61.8%, and 76.5% of the British population aged between 40 and 70 in 2011 had these qualifications respectively.(*23*) See **Table 1** for a description of the participants included in this study. We used inverse probability weights to correct for this selection.

### 3.2 Testing the relevance assumption

Participants born after August 1957, who were affected by the raising of the school leaving age, were 26.6 (95% confidence interval (95%CI): 21.7 to 24.4) percentage points more likely to remain in school after age 15. We used the 74 genetic variants detected in the educational attainment GWAS to construct a weighted genetic score in the UK Biobank. Each variant was weighted by its association with educational attainment in the discovery sample of the GWAS. The educational attainment allele score was more weakly associated with educational attainment. A unit increase in the score was associated with 1.48 additional years of education (95%CI: 1.39 to 1.57) as defined by the International Standard Classification of Education (ISCED). Thus the educational attainment allele score was a strong instrument, but explained less of the variation in educational attainment than the raising of the school leaving age. Neither proposed instrument are likely to suffer from weak instrument bias. The policy reform induced fewer individuals to leave school before the age of 16 (**Figure 1**, top). Whereas the educational attainment allele score was associated with an increased likelihood of remaining in school at all ages (**Figure 1**, bottom).

**Figure 1:**
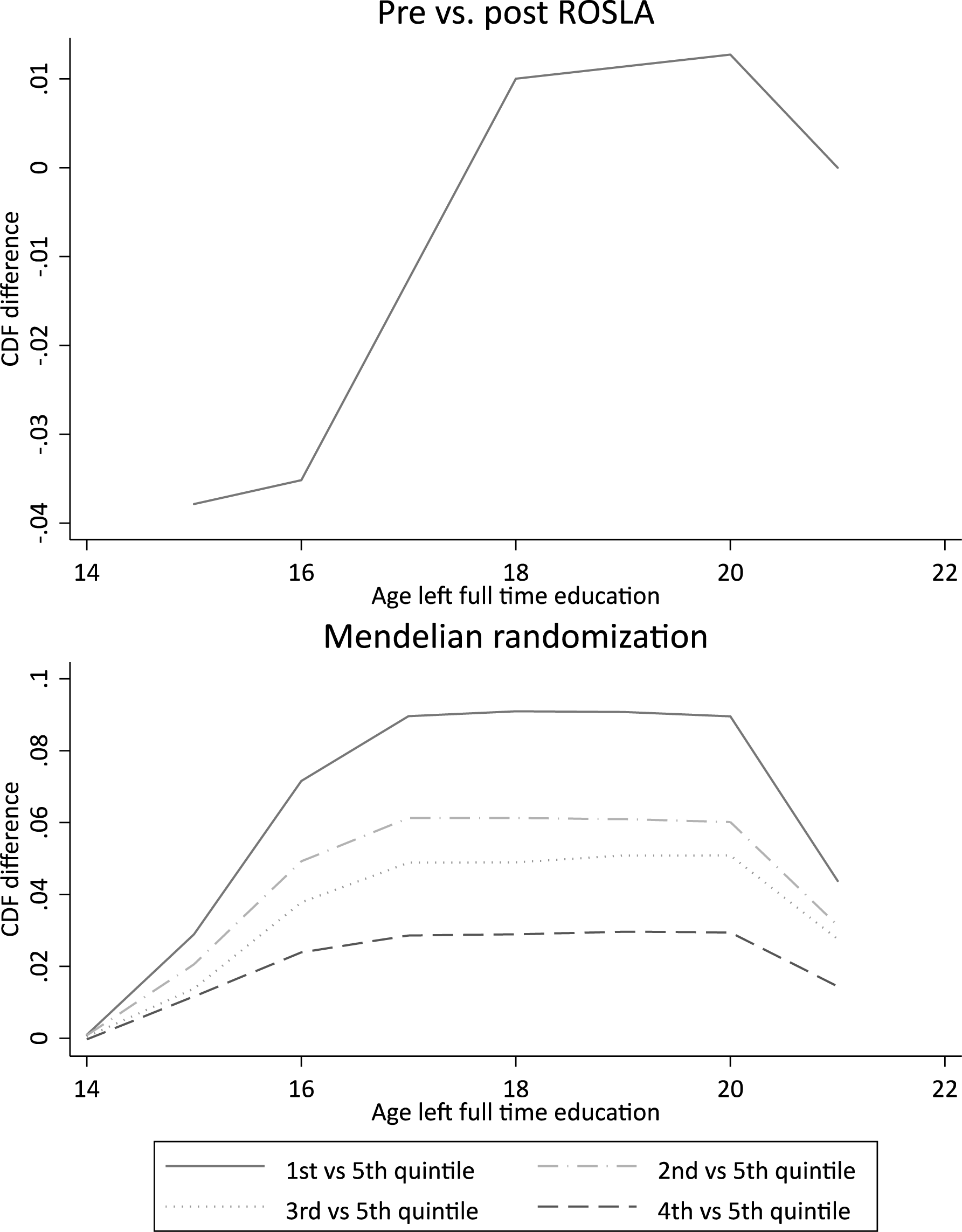
Differences in the age participants left school across the raising of the school leaving age (top) and quintiles of the educational attainment genetic score (bottom). The policy reform substantially reduced the proportion who left school before the age of 16. The genetic scores are associated with a higher probability of remaining in schools at all ages.

### 3.3 Bias component plots

We were concerned that our results may be affected by selection bias or residual confounding. If there was strong selection into the study then this could induce correlations between the instruments and outcomes that are independent in the population. We evaluated this using bias component plots.(*24*) Bias component plots compare the relative bias of the instrumental variable and conventional estimators if an observed covariate was omitted. We assessed the bias associated with 14 non-genetic phenotypes and polygenic scores for 45 traits. The biases for the educational attainment genetic score were similar in size to those for the raising of the school leaving age (**Figures 2 and 3**).

**Figure 2:**
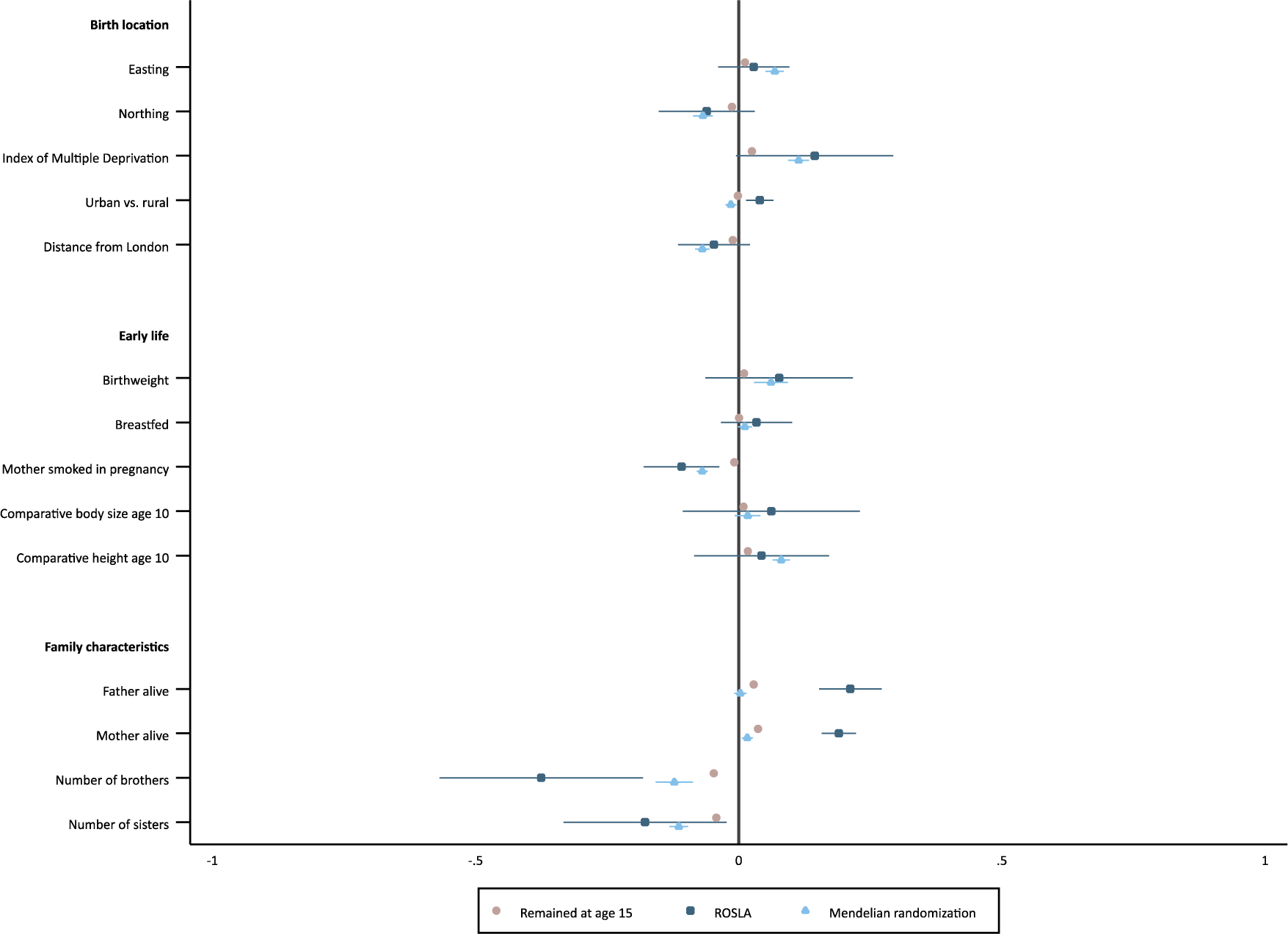
Bias component plots comparing actual educational attainment (ISCED) •, raising of the school leaving age ∎, and the educational attainment genetic score ▴ with non-genetic covariates. There was little evidence that the educational attainment allele score was more strongly associated with the phenotypic covariates than the raising of the school leaving age. There was some evidence that the score was associated with geographic location, but the size of these associations was modest. Notes: Adjusted for month of birth, sex, and the ten principal components of population stratification. Confidence intervals allowing for clustering by month of birth reported. Sample weighted to adjust for under sampling of less educated.

**Figure 3:**
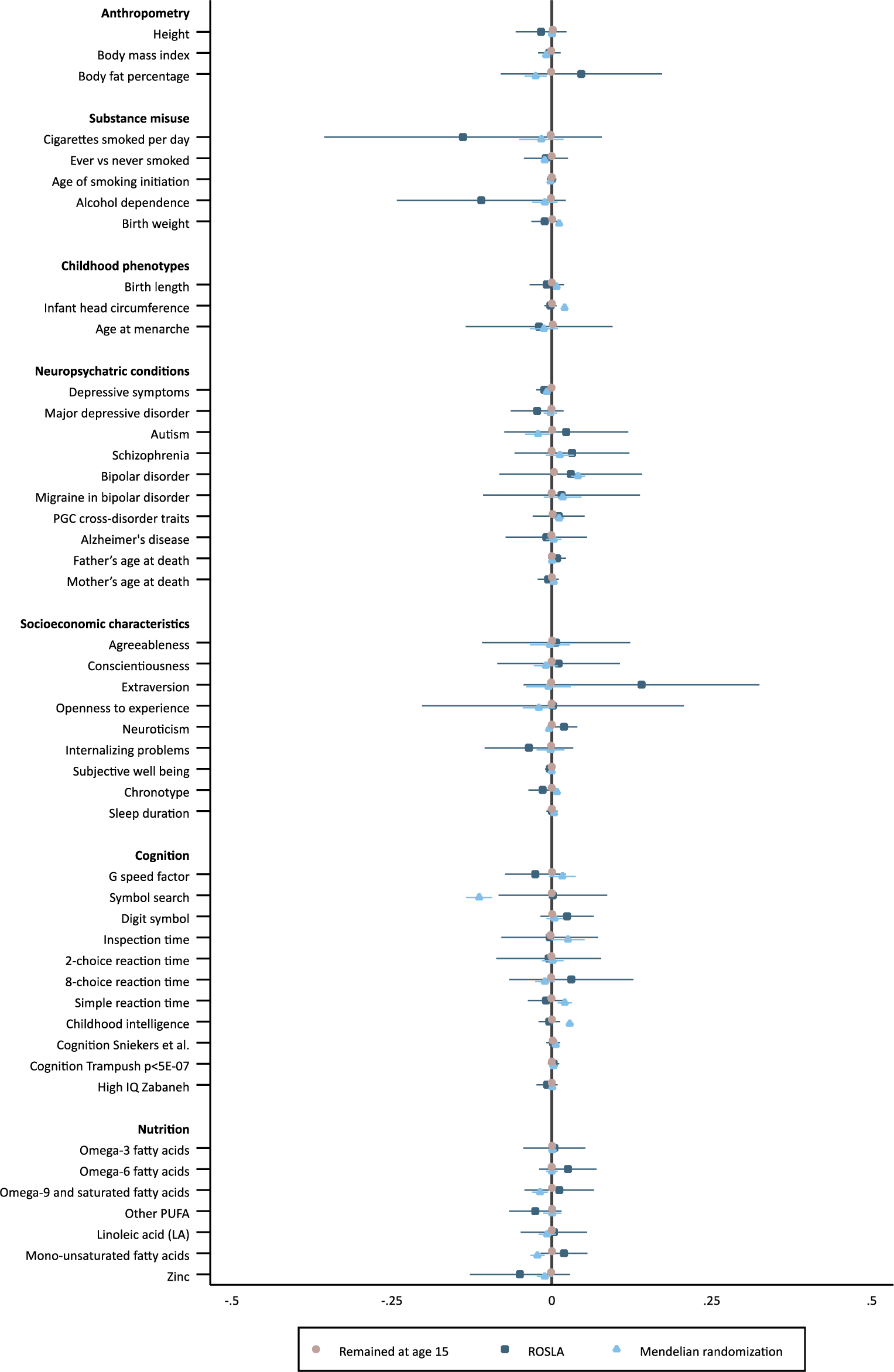
Bias component plots comparing educational attainment (ISCED)•, raising of the school leaving age ∎, and the educational attainment genetic score ▴ with genetic covariates. There was little evidence that the educational allele score was more strongly associated with the genetic confounders than the raising of the school leaving age. Notes: Adjusted for month of birth, sex, and the ten principal components of population stratification. Confidence intervals allowing for clustering by month of birth reported. Sample weighted to adjust for under sampling of less educated.

#### 3.3.1 Phenotypic confounders

The parents of participants affected by the raising of the school leaving age were less likely to have died. These differences are likely to be due to cohort effects. On average participants affected by the reform were one year younger than those who were not affected. Offspring educational attainment may also affect parental mortality.(*25*–*27*) There was little evidence that the reform affected any of the other baseline and childhood phenotypes. There was evidence that the educational attainment genetic score was non-randomly distributed across the UK (**Figure 2**). On average, genetic variants associated with educational attainment were more common in the east and south of the UK. However, the magnitude of these associations was relatively small. There was evidence that the educational attainment genetic score associated with having been breastfed, birthweight, being taller than average at age 10, and whether the participants’ mother smoked in pregnancy. These associations may be driven by dynastic effects or assortative mating. Dynastic effects could occur because on average, participants with more education associated genetic variants will have more educated parents. If more educated parents behave differently, e.g. smoke less in pregnancy, then this could cause an association with the educational attainment genetic score. Assortative mating could induce associations if for example, on average more educated parents choose taller spouses. Nevertheless, these covariates weakly associate with the outcomes. There is little evidence that the covariates associate more strongly with the educational attainment genetic score than the reform. As a result, for many outcomes the bias induced by these covariates is small, for more details see the full adjusted sensitivity analyses below. This suggests that residual confounding due to phenotypic covariates is unlikely.

#### 3.3.2 Genetic confounders

The educational attainment genetic score weakly associated with polygenic scores for other phenotypes including bipolar disorder, childhood intelligence, inspection time, simple reaction time and infant head circumference (**Figure 3**). However, there was little evidence that bias components for the educational attainment genetic score were larger than those for the raising of the school leaving age. This suggests that genotypic confounding is limited.

### 3.4 Effect of educational attainment on outcomes

**Figure 4** plots the estimated effects of an additional year of education on each of the 25 outcomes.

**Figure 4:**
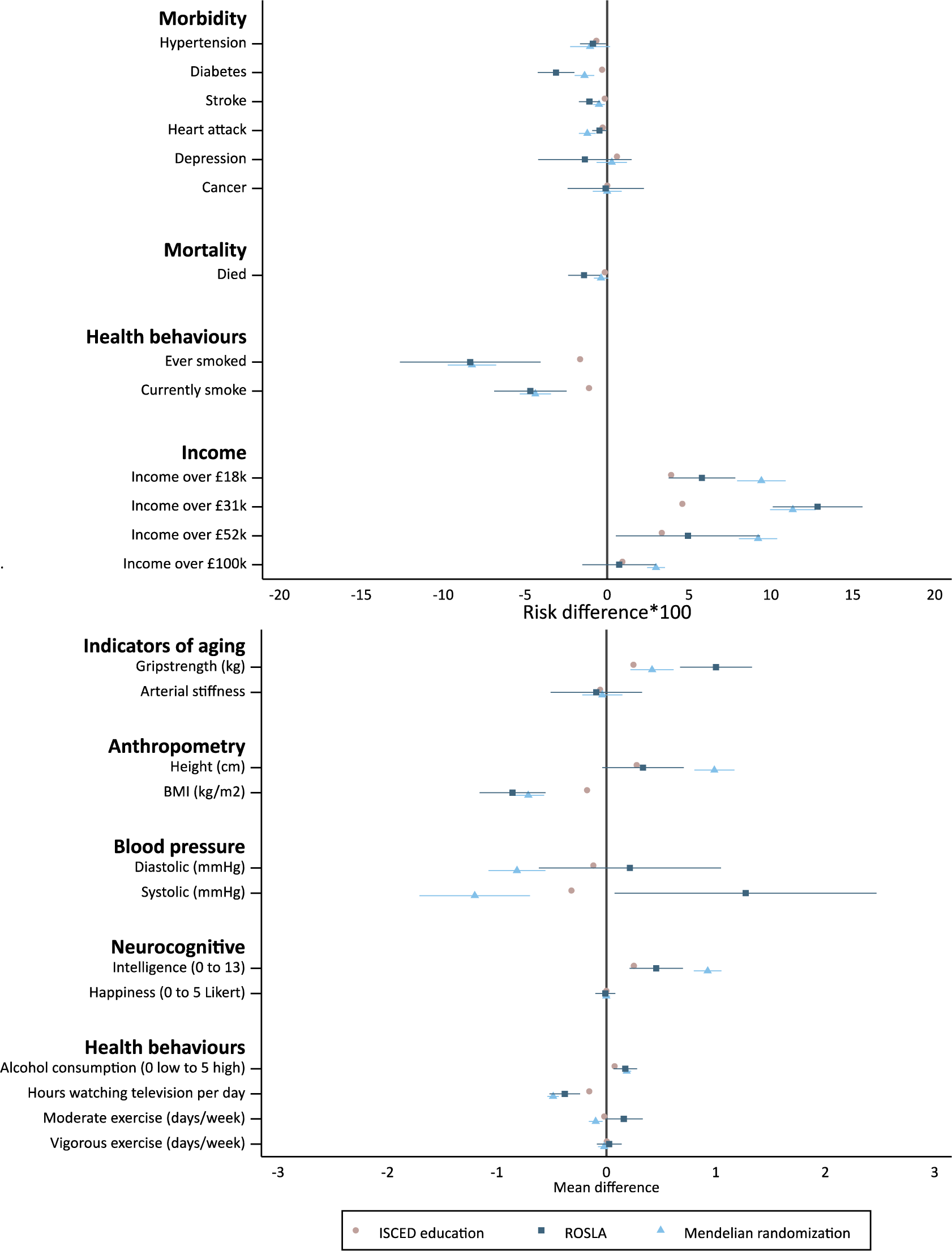
The effect of one additional year of schooling on morbidity, mortality and socioeconomic outcomes, estimated via multivariable adjusted regression ∎, and using instrumental variables raising of the school leaving age ∎, and the educational attainment genetic score Δ The results using both the educational allele score ▴ and the raising of the school leaving age were similar, and suggested that the observational difference is likely to underestimate the difference in outcomes caused by education. Notes: Adjusted for month and year of birth, sex, and the ten principal components of population stratification. Confidence intervals allowing for clustering by month of birth reported. Sample weighted to adjust for under sampling of less educated.

#### 3.4.1 Mortality

Each additional year of education was observationally associated with −0.14 (95%CI: −0.16 to −0.11) percentage points lower mortality. The Mendelian randomization estimates were similar to this but less precise (−0.37 95%CI: −0.80 to 0.06). This effect is larger than the observational association of educational attainment and mortality, but smaller than the effect of remaining in school estimated by the raising of the school leaving age (**Figure 4**).

#### 3.4.2 Morbidity

Observationally, an additional year of education generally associated with improved health. Each year of education was associated with 0.65 per 100 (95%CI: 0.58 to 0.72) fewer cases of hypertension, 0.30 (95%CI: 0.27 to 0.34) fewer diagnoses of diabetes, 0.14 (0.12 to 0.17) fewer strokes, 0.27 (95%CI: 0.24 to 0.30) fewer heart attacks, and 0.60 (95%CI: 0.55 to 0.66) more episodes of depression. There was little evidence of differences in rates of cancer diagnoses. The Mendelian randomization estimates suggested that each year of education reduced the likelihood of being diagnosed with hypertension by 1.04 per 100 (95%CI: −0.18 to 2.25), diabetes by 1.38 (95%CI: 0.78 to 1.97), stroke by 0.50 (95%CI: 0.14 to 0.86), heart attack by 1.21 (95%CI: 0.70 to 1.71). However, the Mendelian randomization estimates provided little evidence of an effect on depression or cancer. The estimates based on the raising of the school leaving age were in the same direction as the Mendelian randomization results. The policy reform suggested larger effects on diabetes and stroke.

#### 3.4.3 Health behaviours

An additional year of education was associated with 1.65 per 100 (95%CI: 1.57 to 1.73) and 1.10 (95%CI: 1.04 to 1.17) fewer ever and current smokers. The Mendelian randomization analysis suggested that the causal effects of education on smoking were substantially larger, 8.25 (95%CI: 6.78 to 9.73) and 4.38 (95%CI: 3.43 to 5.34) fewer smokers per 100. The estimates based on the raising of the school leaving age were similar to those using Mendelian randomization. Each year of education was associated with a 0.07 (95%CI: 0.07 to 0.08) units increase in alcohol consumption. The Mendelian randomization estimates implied the causal effect of an additional year of schooling was 0.19 (95%CI: 0.15 to 0.23). Each year of education was associated with watching 0.16 (95%CI: 0.15 to 0.16) fewer hours of television per day. The Mendelian randomization suggests that this is likely to underestimate the causal effects (0.49, 95%CI: 0.44 to 0.54). A year of education was associated with 0.02 (95%CI: 0.02 to 0.02) fewer days per week of moderate exercise. The Mendelian randomization estimate suggested this underestimated the causal effect (0.10, 95%CI: 0.04 to 0.16). There were only very small associations between educational attainment and vigorous exercise which were similar to the Mendelian randomization and policy reform estimates.

#### 3.4.4 Income

Each additional year of education was associated with a higher probability of having an income above £18,000, £31,000, £52,000 and £100,000 of 3.91 (95%CI: 3.74 to 4.06), 4.59 (95%CI: 4.51 to 4.67), 3.34 (95%CI: 3.20 to 3.48), 0.94 (95%CI: 0.88 to 1.00) per 100 participants respectively. The Mendelian randomization estimates were larger, suggesting 9.42 (95%CI: 7.93 to 10.90), 11.33 (95%CI: 9.94 to 12.72), 9.22 (95%CI: 8.06 to 10.38), and 2.98 (95%CI: 2.44 to 3.53) increase per 100 participants. The raising of the school leaving age analysis were similar in direction and magnitude as the Mendelian randomization estimates but provided little evidence that education affected the probability of having the highest income.

#### 3.4.5 Indicators of ageing

Each year of education was associated with 0.25 (95%CI: 0.24 to 0.26) stronger grips. The Mendelian randomization estimates suggest a larger causal effect of 0.42 (95%CI: 0.22 to 0.61). Education was also associated with lower arterial stiffness 0.06 (95%CI: 0.04 to 0.07). The Mendelian randomization estimate was imprecise, but in the same direction, implying each year of education reduced arterial stiffness by 0.04 (95%CI: −0.14 to 0.22). The estimates based on the raising of the school leaving age suggested a larger effect on grip strength, but similar equivocal effects on arterial stiffness.

#### 3.4.6 Anthropometry

Each additional year of education was observationally associated with a 0.28 (95%CI: 0.27 to 0.29) cm increase in height and 0.18 (95%CI: 0.17 to 0.18) kg/m2 reduction in BMI. The Mendelian randomization estimates suggested larger causal effects of education of 0.99 (95%CI: 0.80 to 1.17) cm increase in height and a 0.71 (95%CI: 0.57 to 0.86) kg/m2 reduction in BMI. The estimated effect on height using the raising of the school leaving age was very similar to the observational association. Whereas the effects on BMI estimated using the reform were much larger than the observational associations, and very similar to the Mendelian randomization estimates. The effect on education on height is likely to be due to pleiotropic or residual population stratification. We investigated this using a negative control outcome: whether the participant reported being taller than average at age 10. Mendelian randomization implied that each additional year of education was associated with being 4.14 (95%CI: 3.03 to 5.24) percentage points more likely to report being taller than average at age 10. We investigated this finding further in the pleiotropy robust sensitivity analyses below.

#### 3.4.7 Blood pressure

Each additional year education was associated with lower diastolic and systolic blood pressure (0.12 mmHg, 95%CI: 0.10 to 0.14 and 0.32 mmHg 95%CI: 0.29 to 0.35 respectively). The genetic analysis suggested the causal effects were in the same direction, but larger (0.82 mmHg 95%CI: 0.59 to 1.08 and 1.20 mmHg 95%CI: 0.70 to 1.71 respectively). There was little evidence the reform affected diastolic blood pressure, and some evidence that it increased systolic blood pressure, however, these estimates are likely to be biased due to age effects.(21)

#### 3.4.8 Neurocognitive

Each year of education was associated with 0.25 (95%CI: 0.25 to 0.26) additional correct answers on the intelligence test, but there was little difference in subjective well-being. The Mendelian randomization estimates suggested that educational attainment caused a 0.93 (95CI: 0.80 to 1.05) additional correct answers, but found little detectable effect on subjective well-being. The estimates of the effect on intelligence based on the raising of the school leaving age were also positive, but were slightly smaller. There was little evidence that the reform affected subjective well-being.

### 3.5 Sensitivity analyses

#### 3.5.1 Weighting to account for non-random sampling

Reanalysing the data without applying inverse probability weights did not materially influence the Mendelian randomization estimates, see **Supplementary Figure 2.**

#### 3.5.2 Association between the educational attainment genetic score and the outcome, the “reduced form”

We present the associations between the educational attainment genetic score and each of the 25 outcomes in **Supplementary Figure 4.** These associations are consistent in direction with the main instrumental variable results presented above.

#### 3.5.3 Robustness of results to adjustment

We investigated whether the results were affected by removing the covariates, including sex, year and month of birth, and the first ten principal components of population stratification. The only estimates that was affected by this were grip strength and height, which attenuated towards the null. See **Supplementary Figure 5** for details. There was little detectable impact of adjusting for a range of confounders, see **Supplementary Figure 6.** The estimated effect of height attenuated modestly. These sensitivity analysis suggests that residual confounding is unlikely to explain our results.

#### 3.5.4 Pleiotropy robust methods

We investigated whether the results could be explained by pleiotropy using MR-Egger, weighted median and weighted mode approaches (**Supplementary Figure 7**). MR-Egger was highly imprecise for all outcomes and provided few inferences. The different estimators provided consistent evidence of causal effects for some of the outcomes including diabetes, Heart attack, mortality, smoking, income, grip strength, BMI, blood pressure, intelligence, alcohol consumption, and exercise. There was evidence of differences in the estimates for height, which may indicate the inverse variance weighted and two-stage least squares estimates above suffer from pleiotropy. The weighted mode estimator suggested little effect of educational attainment on height. We present the I2 statistics of the heterogeneity in estimated effects of education across the 74 genetic variants in **Supplementary Table 2.**

## 4 Discussion

Our findings suggest that the differences in many later life outcomes between educational groups are likely to be caused by education. There was evidence that genotypic perturbations in educational attainment associated with morbidity, including the risk of hypertension, diabetes, stroke, heart attack, and mortality. Furthermore, these results imply that education reduces the risk of currently or ever smoking, increases household income, lowers blood pressure and increases scores on intelligence tests. However, there was evidence that education reduced rates of moderate exercise and increased alcohol consumption. Our sensitivity analyses suggest that confounding by genotypic or phenotypic confounders, or specific forms of pleiotropy are unlikely to explain our results.

Triangulating across multiple sources of evidence can help provide stronger evidence of causal effects.(*28*) Here, we found that the two natural experiments gave remarkably similar results. The two sources of variation, Mendelian randomization and the raising of the school leaving age, have distinct causes of and direction of bias. The similarity in results strengthens the case that education has causal effects. The raising of the school leaving age affected relatively low-ability students, who were forced to remain in school for an additional year.(*1*) In contrast, our Mendelian randomization results exploit variation across the entire distribution of educational attainment, estimating an average effect of an additional year of schooling for everyone from those who leave school at 15 to graduates (see **Figure 1**). A priori there was little reason to assume that a year of additional schooling will have the same effects on a high school leaver as on a graduate. Surprisingly we found relatively little evidence that educational attainment had heterogeneous effects. The estimates from the two natural experiments are remarkably similar, both in direction and in many cases magnitude. There was very little evidence of heterogeneity in the effects identified by different variants. The effect of an additional year of education on smoking is comparable to other studies using natural experiments. For example, Grimard and Parent (2007) used data from the US Current Population Survey and the Vietnam draft to estimate that in 1995-99 an additional year of schooling caused a 7.97 (95%CI: 3.15 to 12.79) and 11.13 (95%CI: 5.54 to 16.72) percentage point reduction in probably of currently or ever smoking.(*29*) For other outcomes, such as measured blood pressure the results are very different. The effects of education on blood pressure estimated using the raising of the school leaving age may reflect non-linear cohort effects as previously discussed.(*21*)

We found some evidence that the educational attainment genetic score correlated with baseline covariates, including birth weight, being taller than average at age 10, mother smoked in pregnancy, parental mortality and geography. These associations may reflect dynastic effects or assortative mating (**Figure 3**). If there is assortative mating, then this could induce associations between education variants and variants for other traits. For example, if highly educated people assortatively mate with taller spouses, then the Mendelian randomization estimates of the effect of education on height would be positively biased. These effects may explain the implausible Mendelian randomization estimate of the effect of education on height. A sub-sample (N= 310,230) of the study provided information on whether they were taller than average at age 10, this variable cannot be affected by completed years of educational attainment. When we adjusted for being taller than average at age 10 the estimated effect falls from 0.93 (95CI: 0.78 to 1.09) to 0.62 (95CI: 0.46 to 0.78) cm increase in height per year of education. This result suggests that the effects on height may be induced by assortative mating, dynastic effects or population stratification.

A limitation of our study is that we used a non-representative sample. We have addressed this using inverse probability weights. The weights made little differences to the Mendelian randomization estimates. This suggests that sample selection bias is unlikely to affect our results.

The Mendelian randomization estimates can suffer from bias due to assortative mating or dynastic effects, however except height, adjusting for measured baseline covariates had little affect on our results (**Supplementary Figure 5** and **6**). Our results could reflect either direct effects of the participants’ educational attainment, assortative mating between their parents, dynastic effects of their parents’ education or differences in ancestry not accounted for by the principal components (**Supplementary Figure 1**). These potential explanations could be evaluated using either offspring-mother-father trios or sibling designs. Okbay and colleagues (2016) found little evidence that the effects of the genome-wide significant education variants attenuated after controlling for family structure.(*30*) However, these analyses may not have had sufficient power to detect dynastic effects. Kong and colleagues investigated this using a sample of parent and offspring from Iceland.(*31*) They found that a polygenic score for education, made up of alleles that were not inherited, was associated with offspring’s education. The association with the non-inherited polygenic score was 29% of the size of the association with the inherited genetic score. This suggests that the effects we identify are likely to represent a combination of the effect of the participants’ education and their parents’ education. Kong and colleagues results provide an upper bound for the contribution of parents’ education of 29%. The contribution of parents’ education to our results will be smaller if the direct effect of parents’ education on each outcome is smaller than the effect of the participant’s own education. To determine the relative contributions of parent versus offspring education will require large samples of parent-offspring data.

A further limitation is that the genetic variants may have pleiotropic, or direct effects on the outcomes, as their biological mechanisms of effect are unknown. However, our estimates were similar when using the weighted median and mode estimates. An exception to this was height, where the mode and median based estimate suggested smaller effects.

In conclusion, two independent natural experiments suggest that education has wide-ranging effects on important outcomes measured much later in life. Importantly, the two experiments affected different educational choices – one exclusively affecting those at the bottom of the distribution, the other affected education levels across the whole distribution – and yet find effects of a similar magnitude. This suggests a common treatment effect of additional education on many health behaviours and outcomes.

## Supplementary Materials

### 6 Materials and Methods

#### 6.1 Data

The UK Biobank sampled 503,325 people via 23 study centres in urban areas across the United Kingdom. The study invited people between the ages of 40 and 70 to attend an assessment clinic between 2006 and 2010. The participants completed surveys, and had detailed phenotypic measurements and provided samples of blood.

#### 6.2 Genotyping

DNA was extracted from the blood sample which was genotyped using UK BiLEVE Axiom and UK Biobank Axiom arrays. Full details of the genotyping and imputation procedure are available.(*32*) The genotyping data quality control process consisted of the following steps. First SNPs were set to missing when there was evidence of clustering across chips, batch or plate effects, departure from Hardy Weinberg equilibrium, sex effects, array effects or discordance across the genotyping controls (at p<10-12). On average these exclusion criteria affected 7,704 (0.01%) of SNPs per batch. Second, 968 participants who had less than 5% call rates or extreme heterozygosity were excluded. Third, individuals from a European ancestry were identified by projecting each participant onto the principal components from the 1000 genomes project. Fourth, we excluded participants who were as genetically alike as third-degree relatives or more. Fifth, we excluded participants whose self-reported gender did not match their genetic sex.

#### 6.3 Educational attainment

For the observational and Mendelian randomization analyses, we used years of schooling as the exposure, for the raising of the school leaving age we used whether someone had remained in school after the age of 15. Within the instrumental variables framework, this means that both exposures indicate the effect of an additional year of schooling. We derived each participants’ years of schooling using the information they provided using a touch screen survey as part of their assessment centre visit. This survey included questions on the educational qualification the participant had (i.e. did they have a degree, A-levels). We used these variables to derive a measure of educational attainment based on the International Standardized Classification of Education (ISCED). This definition was used by Okbay and colleagues, we recoded the number of years of education each category referred to be consistent with the UK education system. See the supplementary materials for a detailed coding. If the participant stated they did not have a degree they were asked at which age they left school. We used these survey responses to derive an indicator of whether the participant remained in school after age 15. If the participant did not have a degree, then this was equal to one if they stated they left school after the age of 15, otherwise, it was set equal to zero. If they stated they had a degree, then this variable was set to one.

#### 6.4 Outcomes

##### 6.4.1 Morbidity

The participants completed questionnaires about whether a doctor had diagnosed them with high blood pressure, stroke, or a heart attack. They were asked if they had been diagnosed with diabetes. We set this outcome to missing if they received a diagnosis before the age of 21. They were also asked if they had experienced episodes of depression. Finally, cancer diagnoses were defined using linked cancer registry data.

##### 6.4.2 Mortality

Mortality was defined using linked NHS mortality records. This dataset included the date of death for all participants which occurred after attending the clinic until 17th of February 2014.

##### 6.4.3 Health behaviours

The participants were asked detailed questions about their smoking history. From this information was derived about whether they currently or had ever smoked. They were asked about their alcohol consumption, which was coded as an ordinal variable (0=never, 1=special occasions only, 2=one to three times a month, 3=once or twice a week, 4=three or four times a week, and 5=daily or almost daily). They were asked how many hours they spent watching television per day, and the number of days per week they did 10 minutes or more of moderate or vigorous physical activity.

##### 6.4.4 Income

The participants were asked their average total before-tax household income. We transformed this into four binary variables indicating whether their income was above £18,000, £31,000, £52,000, or £100,000.

##### 6.4.5 Indicators of ageing

During the clinic visit measures of grip strength were taken on both hands using a Jamar J00105 hydraulic hand dynamometer. The measures for each hand were averaged and residualized to account for between device differences which accounted for 2.92% of the variation in grip strength. Pulsewave arterial stiffness was measured from the finger using an infra-red sensor (PulseTrace PCA2, CareFusion, USA). These measurements were residualized to account for between device differences which account for 2.53% of the variation in arterial stiffness.

##### 6.4.6 Anthropometry

We derived height and BMI using the participant’s standing height was measured using a Seca 202 measuring rod, and their weight was measured.

##### 6.4.7 Blood pressure

The participants’ blood pressure was measured twice using an Omron 705 IT electronic blood pressure monitor. These measurements were averaged to give a measure of diastolic and systolic blood pressure.

##### 6.4.8 Neurocognitive

The participants used a touchscreen to complete a battery of 13 fluid intelligence questions. The participants were given 2 minutes to answer as many questions correctly as possible. The participants were also asked if they were extremely, very, or moderately happy or unhappy.

#### 6.5 Statistical methods

We investigated two potential instrumental variables for educational attainment, 1) the raising of the school leaving age in 1972, 2) an allele score (called the “educational attainment genetic score”) constructed using results from the discovery sample of a GWAS of years of education in an independent sample.

##### 6.5.1 The raising of the school leaving age

In September 1972, the minimum school leaving age in the United Kingdom increased from 15 to 16. This forced participants who would otherwise have left school to remain in school for an extra year. This policy reform has been used by a large number of research papers to estimate the effect of schooling on later outcomes. It is a plausible natural experiment because parents of children affected by the reform could not have anticipated the change in the law at the time of conception. So on average individuals affected by the reform will be similar to those who were not affected. We used a 12-month bandwidth. This analysis compares the outcomes of the first individuals in the first school cohort affected by the reform to the outcomes of the last cohort who were not affected. We accounted for linear secular trends in the outcome using a difference in difference design. We subtracted the average year-on-year difference for the cohorts born in the ten years before and after the reform. See Davies and colleagues for further details.(*21*)

##### 6.5.2 Allele scores for educational attainment

We constructed the allele scores using the 74 SNPs which were associated with years of education (p<5e-08) in the discovery sample of the educational attainment GWAS.(*30*) Five SNPs reported by the GWAS were not available in the Haplotype Reference Consortium (HRC) panel, we replaced these SNPs with proxies that were in perfect LD and the HRC panel. See **Supplementary Table 1** for a list of the specific GWAS used. The allele scores are the weighted sum of the number of education increasing alleles for each participant. The contribution of each SNP to the score was weighted by the size of the coefficient reported by the GWAS. The effect alleles of all GWAS results were harmonised to be consistent with the UK Biobank genome-wide data. We excluded palindromic SNPs with an allele frequency of 0.3 or above. We checked for consistency between the effect allele frequency between the GWAS and UK Biobank data. The allele frequencies were correlated 0.9913, and the maximum difference in allele frequency was 0.091.

#### 6.6 Specification tests

Instrumental variables are defined by three assumptions, 1) they must be associated with the risk factor of interest, 2) they must have no common cause with the outcome (no confounding), and 3) they must have no direct effect on the outcome (the exclusion restriction). We tested whether the first assumption held using a partial F-statistic of the association of the instrument and the exposure. We investigated the plausibility of the second assumption by estimating the association of each of the instruments and a broad set of phenotypic and genetic confounders (defined below). We used covariate bias plots to account for the relative strength of the instruments.(*24*) Covariate balance plots are the ratio of the association of the instrument and a measured confounder and the association of the instrument and the exposure (educational attainment).(*33*) We estimated these terms using the generalised method of moments.(*34*)

Covariate balance plots allow for direct comparison of the relative bias caused by omitting each observed covariate from an instrumental variable or multivariable-adjusted regression analysis. The third instrumental variable assumption, that the instrument has no direct effect on the outcome, can potentially be investigated (falsified) if there are measures of alternative mediating pathways. The UK Biobank recruited participants over the age of 40 after they completed full-time education; therefore it has few suitable measures to evaluate this assumption. We evaluated this assumption using a small number of available covariates and a set of genetic covariates.

Previous studies have found that the raising of the school leaving age affected students who would otherwise have left school at the age of 15. These students were generally less academic, and most chose to leave at the new minimum age of 16. This meant the reform had little effect on the proportion of students remaining in school to the age of 18 or attending university. This means the raising of the school leaving age estimates the effect of an additional year of schooling at age 15. In contrast, the genetic variants associated with years of education are likely to associate with educational outcomes across the entire life course, and the probability of remaining in school at age 15, 16, 18 and the likelihood of obtaining a degree. The effect identified by the genetic variants is, therefore, a weighted average of the effects of an additional year of educational attainment across the entire education distribution.(*35, 36*) We investigated this by plotting the difference in the cumulative density of educational attainment by the policy reform and between the quintiles of the educational attainment allele score.

##### 6.6.1 Covariates: Non-genetic

We also assessed the association of the proposed instruments with 14 phenotypes that are measures of circumstances around the time of birth, childhood, or relating to family background. We assessed: birth location (easting, northing, Index of Multiple Deprivation, urban vs. rural and distance from London), early life factors (birth weight, whether the participant was breastfed, or their mother smoked during pregnancy, their comparative body size and height at age 10, and their number of brothers and sisters). Some of these variables occurred after conception. However, they are unlikely to be directly affected by the educational attainment of the participant, so can provide useful evidence of the plausibility of the instrumental variable assumptions.

##### 6.6.2 Covariates: Genetic

Genomes are determined at conception. Mendel’s law of independent assortment states that, in the absence of any other process (e.g. assortative mating, sample selection bias or variants in linkage disequilibrium), genetic variants for one trait will be inherited independently of another. Thus people who have many and few education increasing alleles should, on average, have a similar number of alleles known to increase BMI. However, individual SNPs are unlikely to provide sufficient statistical power. One way to increase the power is to assess this assumption using multiple variants for a given trait combined in polygenic risk scores. For example, if Mendel’s law of independent assortment holds, then we would expect a polygenic risk score for educational attainment to be independent of a polygenic risk score for BMI. If the education polygenic score associates with scores for other traits then it suggests that Mendel law is unlikely to hold.

We evaluated this using genetic scores for 45 traits extracted from MR-Base (**Supplementary Table 1**).(*37*) We constructed the scores from extracted SNPs that were associated with each trait at p<5e-5. We used a lower threshold than is usually used for genome-wide significance (p<5e-8) to define the scores because we wanted to maximise the explanatory power of the scores. Furthermore, it is not possible for the educational attainment genetic score to have pleiotropic effects on the other polygenic scores. We LD pruned the SNPs for each trait using a threshold of r^2^>0.001 across a distance of 10,000kb. We excluded SNPs from these scores that were in LD (r^2^>0.001) with the 74 SNPs identified as associated with educational attainment at the genome-wide level (p<5e-08) the educational attainment GWAS.(*30*) This resulted in a set of SNPs in independent points in the genome for each trait. We constructed allele scores equal to the sum of the effect alleles for each trait.(*38*) The contribution of each SNP to the allele score was weighted by the coefficient reported in the GWAS for that trait. We harmonised the direction of SNP effects between UK Biobank and the GWAS. Finally, we checked for consistency of the allele frequency reported in the GWAS and the UK Biobank data.

#### 6.7 Sample selection

The UK Biobank is a highly non-random sample, that over sampled those with degrees and sampled relatively few people who have little education or no qualifications. This method of sampling could cause collider bias in our sample if the sample selection relates to both the outcome and the potential instruments. We accounted for this non-random sampling using inverse probability weights. Individuals who reported leaving school at age 15 were weighted by 34.29, and the selected participants who left school age 16 or older were weighted by 12.37 (weights rounded to two decimal places).(*39*) We investigated whether our results were sensitive to the specification of these weights in a sensitivity analysis in the supplementary materials.

#### 6.8 Instrumental variable estimators

In our main analyses, we report the estimates of the effect of educational attainment using two-stage least squares for continuous outcomes, and additive structural mean models for binary outcomes.(*40*–*42*) These identify the causal mean and risk differences respectively. The main analyses adjust for sex, age, and month of birth. We report sensitivity analyses without adjustment and also additionally adjusting for the covariates associated with the educational attainment genetic score in **Figure 2** in the supplementary materials.

##### 6.8.1 Identifying assumptions

The two-stage least squares estimates of the effect on the continuous outcomes can be point identified by assuming a constant effect of education on the outcome, i.e. that an additional year of education causes the same unit change in the outcome for everyone. Alternatively, we could assume that the education allele score has a monotonic effect on education. That is, an additional educational attainment associated allele will increase the likelihood of having a higher level of education in everyone. This estimate identifies the “local average treatment effect” (LATE). This parameter is the effect of education on individuals whose educational choices were affected by the score. For the structural mean models for binary outcomes, we can either assume monotonicity, which is interpreted in the same way as above, or that the effects of education are the same regardless of the number of education variants each participant has. These estimates can be interpreted as the effect of education on individuals who chose to receive a given level of education.

##### 6.8.2 Sensitivity analyses

We investigated the raw association of the education score and each of the outcomes, the so-called “reduced form”. The reduced form is a general test of causation under a minimal set of assumptions, i.e. it does not require point identification. Reduced form and instrumental variable estimates should be consistent in direction. We were concerned that our results could suffer from residual confounding given the non-random geographic distribution of alleles. So we investigated whether our results were sensitive to removing the basic controls (sex, year and month of birth, the first ten genetic principal components). We also investigated whether our results were sensitive to including a richer set of controls in the subset of participants with these data.

Our results could be affected by bias due to pleiotropic effects of the variants. That is if the variants directly affected the outcomes via pathways other than education. We investigated this using MR-Egger, weighted median, and weighted mode estimators in summary data analyses.(*43*–*45*) We included the inverse variance weighted estimates for comparison. These estimates use the educational attainment GWAS discovery sample coefficients. We assessed the variability in the estimated effect of education across the 74 genetic variants using the I2 statistic.(*46*)

##### 6.8.3 Data availability

The analytic datasets used in the study have been [reviewer note - will be] archived with the UK Biobank study. Please contact for further information.

##### 6.8.4 Code availability

All analyses were conducted in StataMP 14.0.(*47*) The code used to generate these results have been [reviewer note - will be] archived at (https://github.com/nmdavies/UKbiobank-MR-vs-ROSLA).

---

**Supplementary Figure 1:**
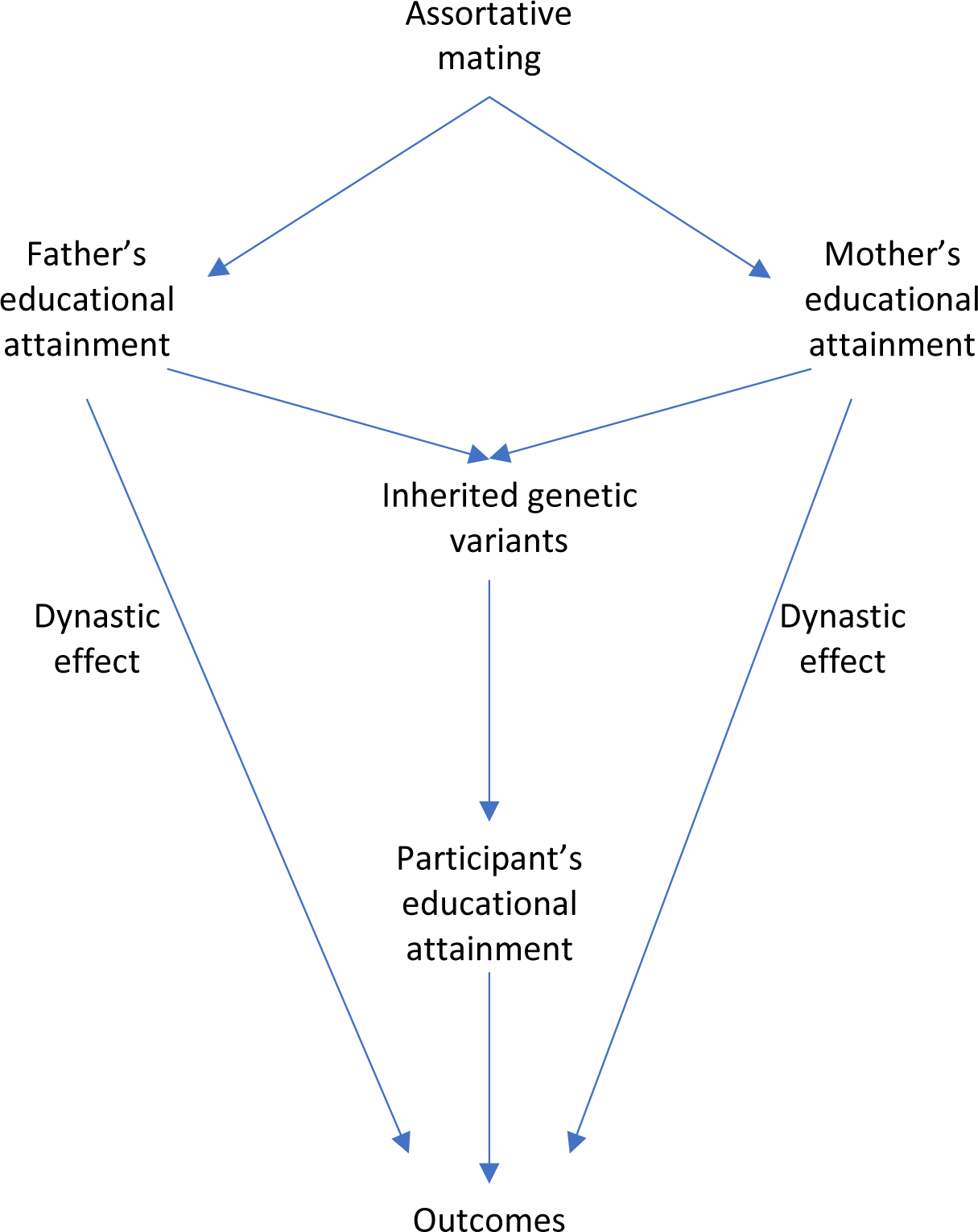
The Mendelian randomization estimates identify the effects of education on the outcomes using directly inherited genetic variants known to associate with educational attainment. These effects could mediated via 1) an effect of the participants’ educational attainment on their later outcomes, 2) a direct effect of parents’ phenotypes on their offspring’s outcomes, 3) or assortative mating on education and other phenotypes (e.g. height) between parents. Dynastic effects would lead to the Mendelian randomization estimates to overestimate the direct effects of educational attainment. Assortative mating would induce associations between the 74 SNP known to associate with education and other variants associated with education or related traits across the genome.

**Supplementary Figure 2:**
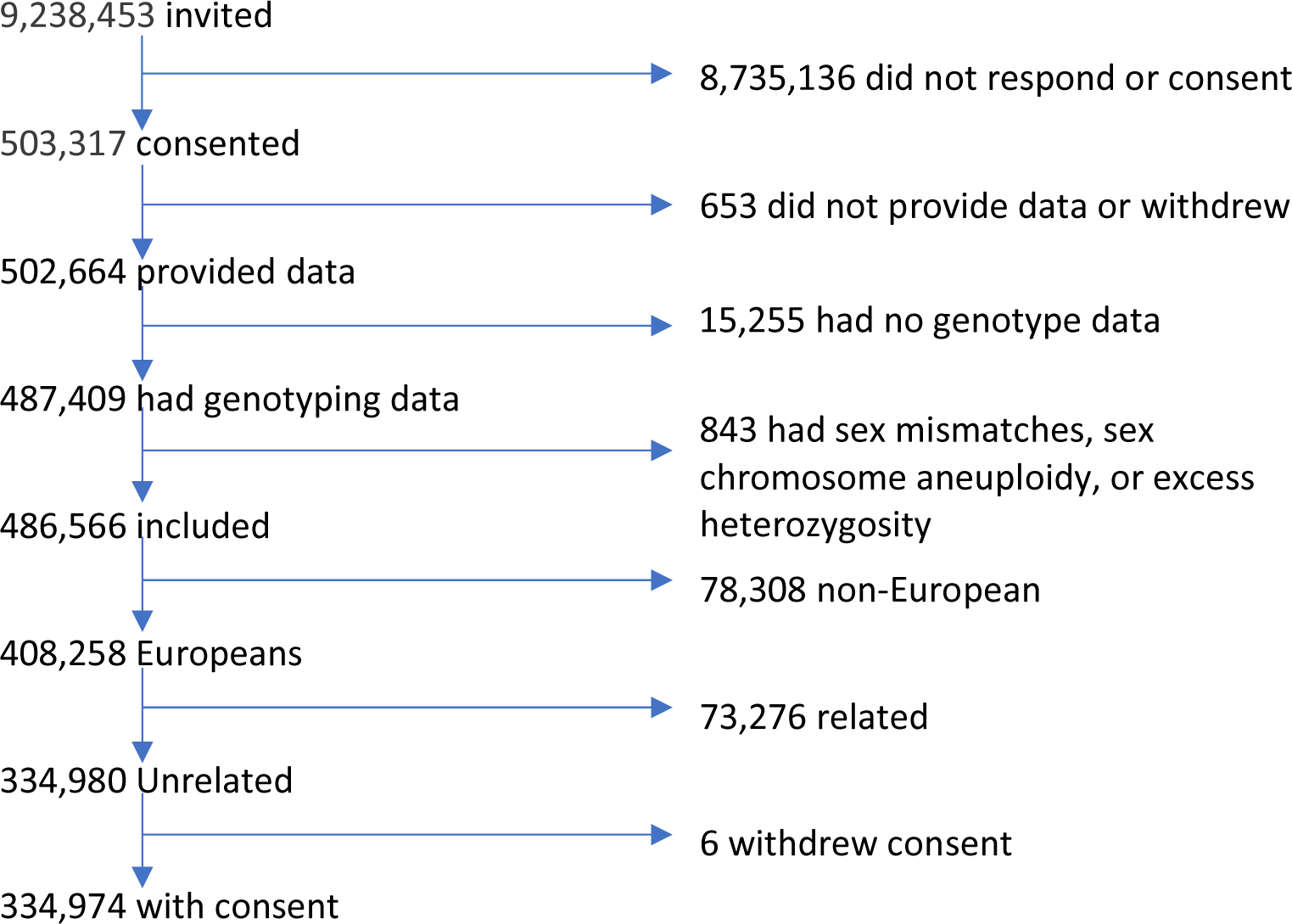
Selection of participants into the study

**Supplementary Figure 3:**
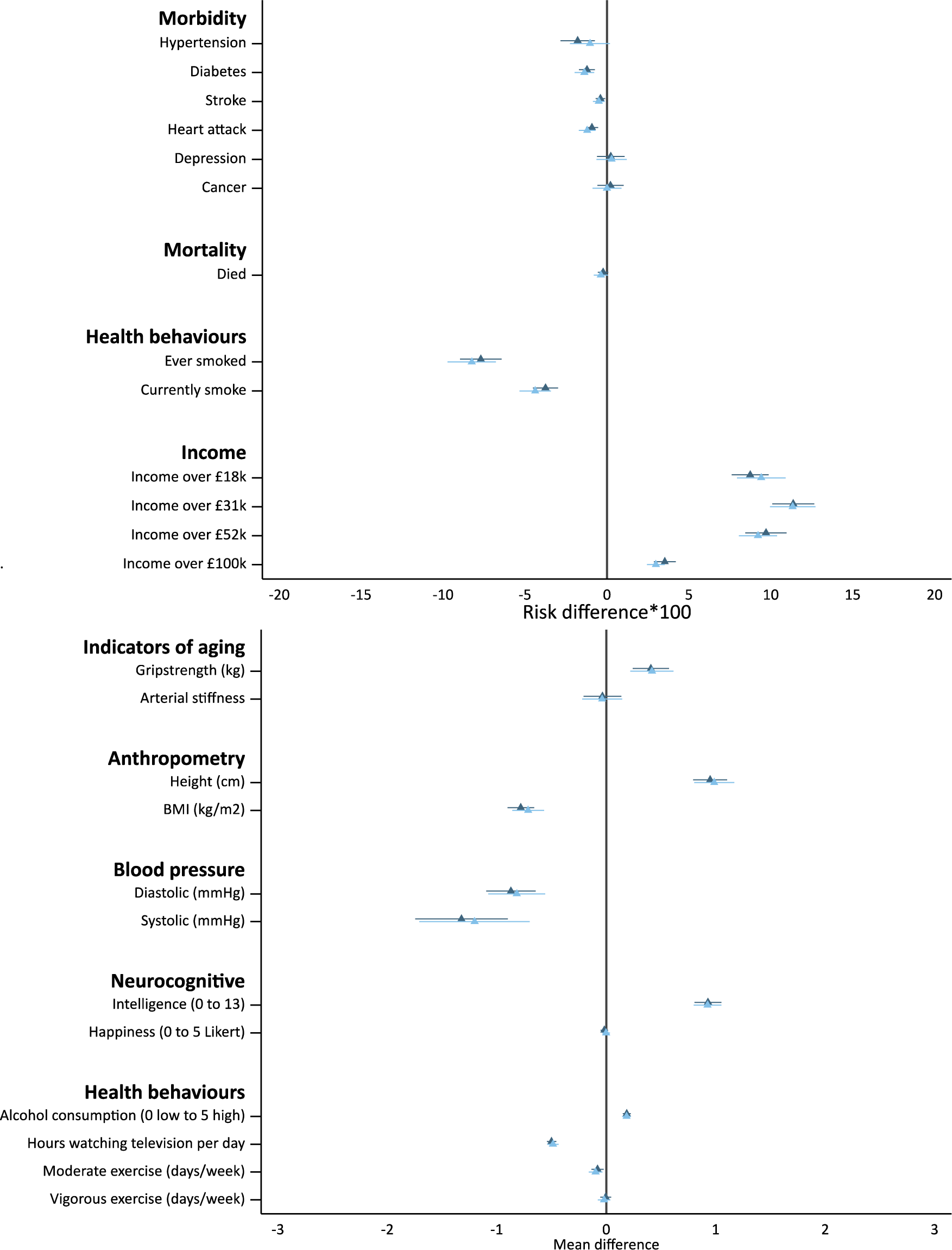
The effect of one additional year of schooling on morbidity, mortality and socioeconomic outcomes estimated using the educational attainment genetic score with and without weighting for under-sampling of less educated ▴ and ▴ respectively. The weighting did not affect the estimates. Notes: Adjusted for month and year of birth, sex, and the ten principal components of population stratification. Confidence intervals allowing for clustering by month of birth reported.

**Supplementary Figure 4:**
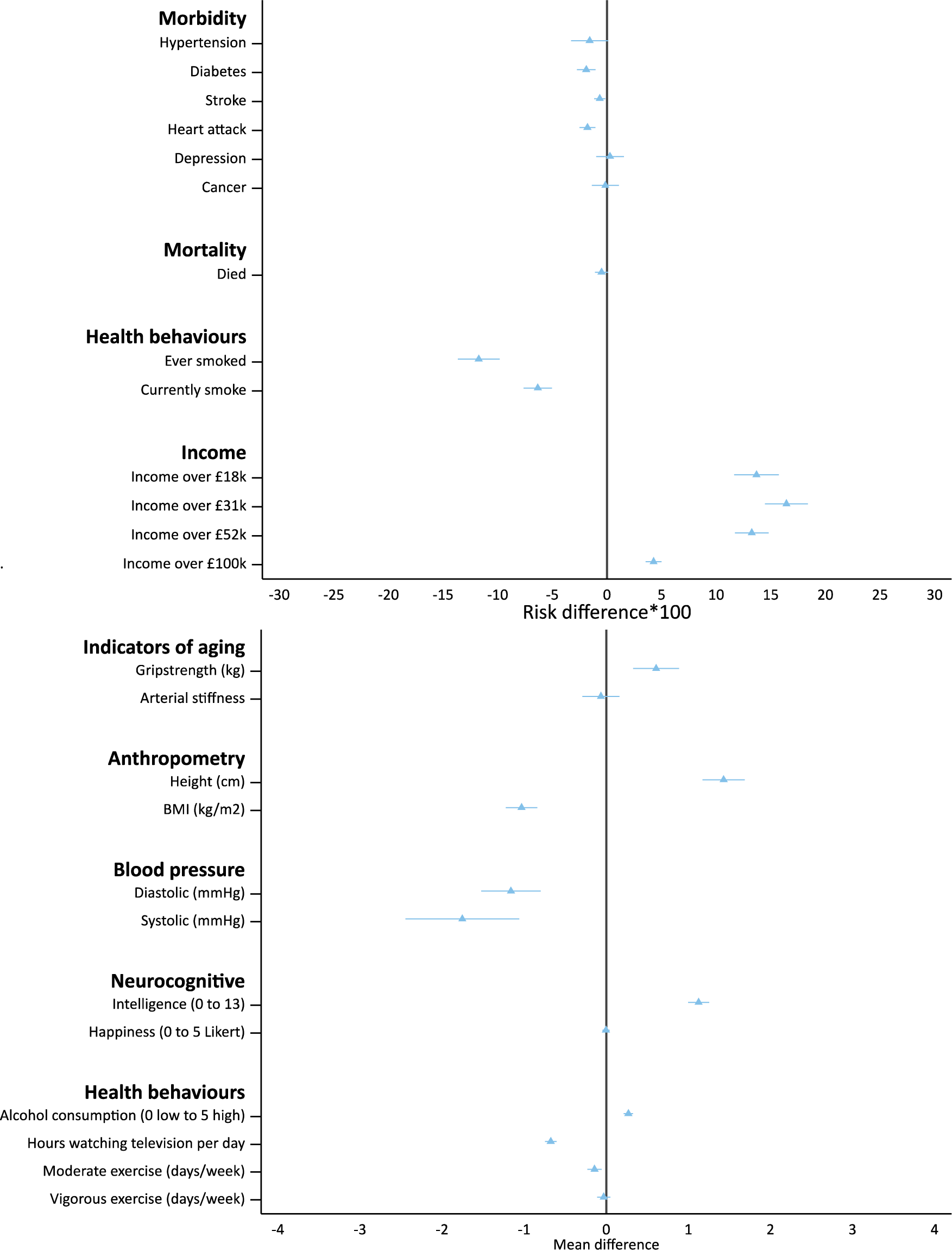
The association of morbidity, mortality and socioeconomic outcomes and the educational attainment genetic score ▴ (the “greduced form”). These estimates are consistent with the main analyses presented in Figure 3. Notes: Adjusted for month and year of birth, sex, and the ten principal components of population stratification. Confidence intervals allowing for clustering by month of birth reported. Sample weighted to adjust for under sampling of less educated.

**Supplementary Figure 5:**
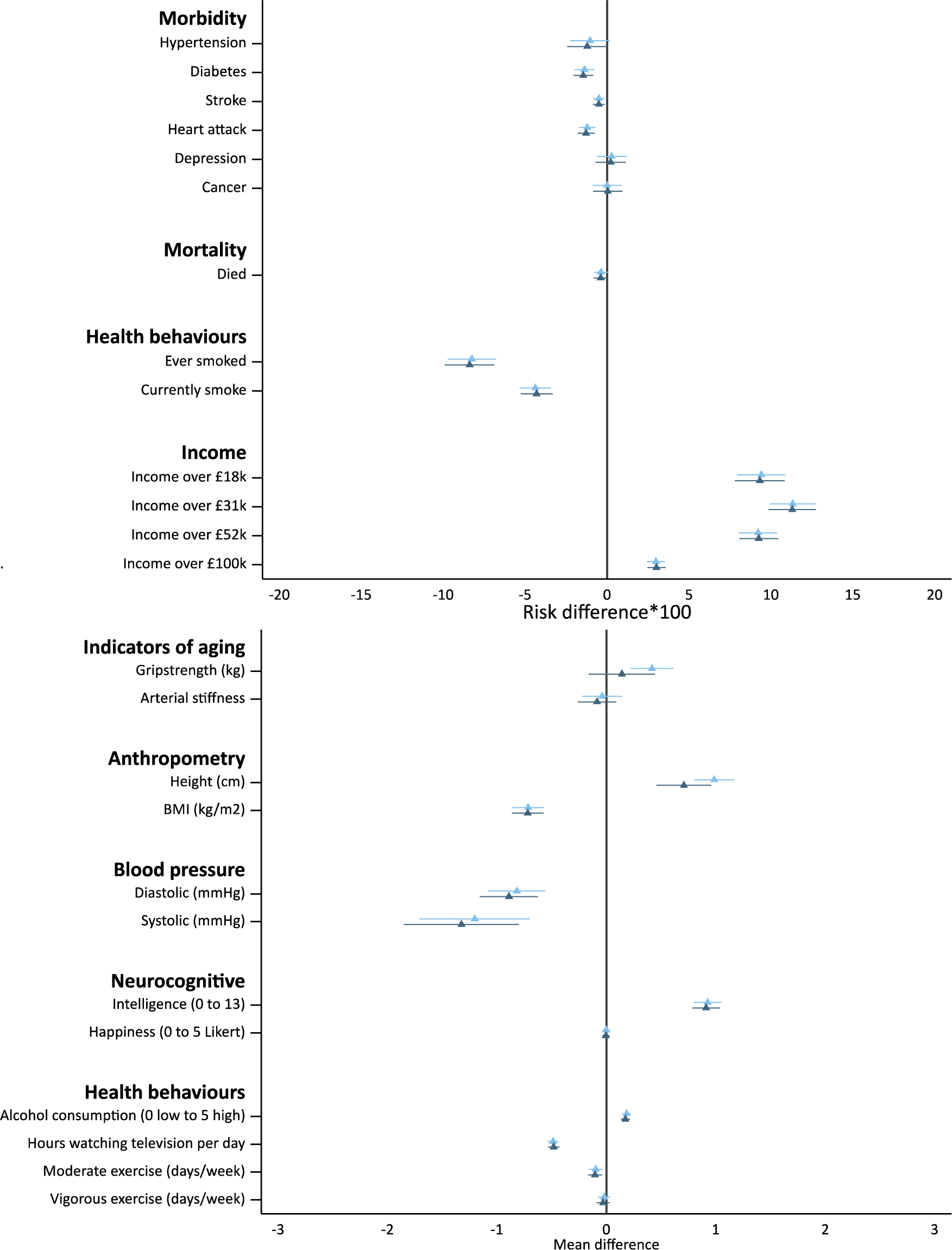
The effect of one additional year of schooling on morbidity, mortality and socioeconomic outcomes estimated using the educational attainment genetic score with and without adjusting for the sex, month and year of birth and principal components of population stratification ▴ and ▴ respectively. The Mendelian randomization estimates were robust. Notes: Confidence intervals allowing for clustering by month of birth reported. Sample weighted to adjust for under sampling of less educated.

**Supplementary Figure 6:**
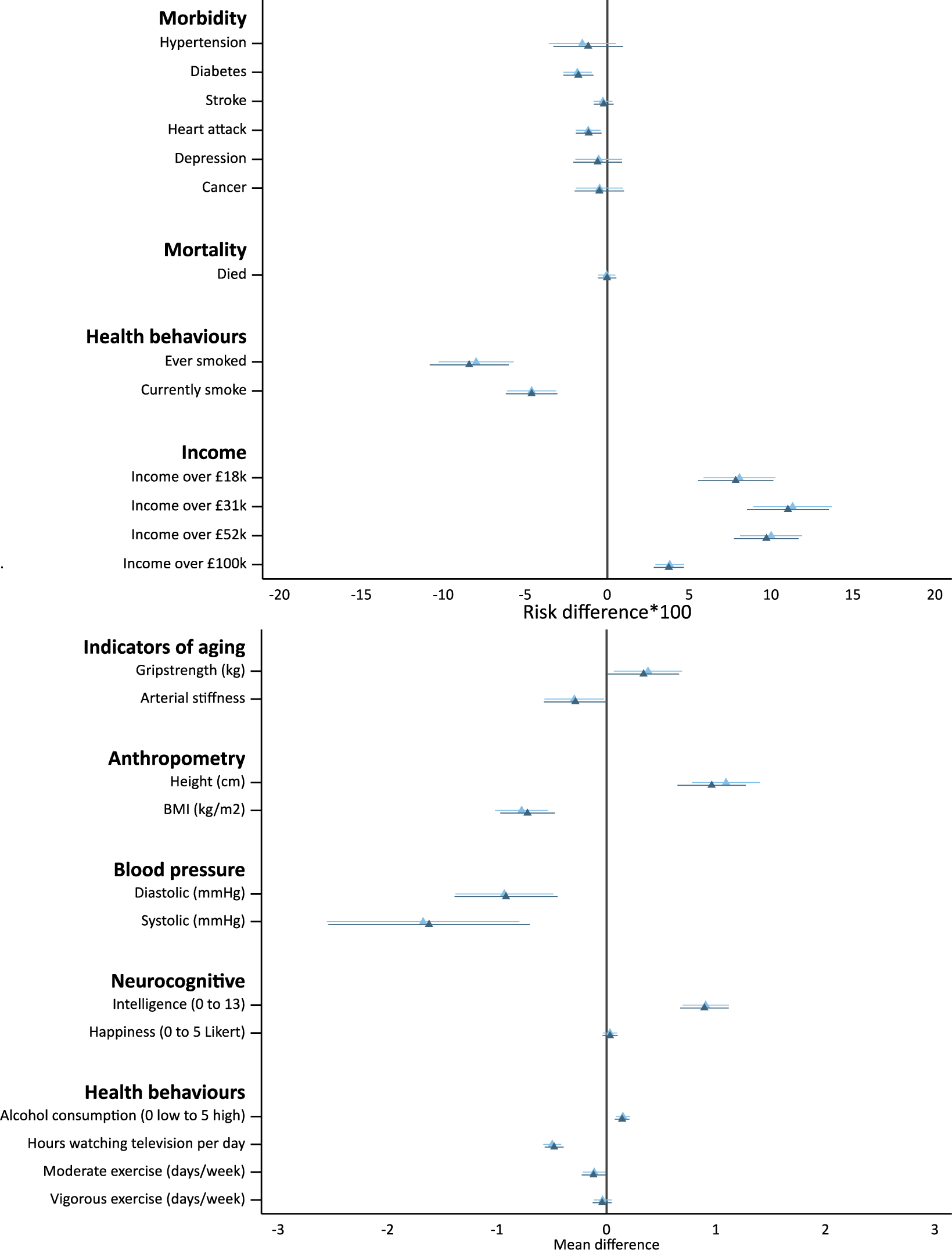
Fully adjusted results. The effect of one additional year of schooling on morbidity, mortality and socioeconomic outcomes estimated using the educational attainment genetic score with and without additionally adjusting for breastfeeding, mother smoked during pregnancy, birth weight, birth location and deprivation (easting, northing, and distance to London) ▴ and ▴ respectively. The fully adjusted estimates were comparable after including additional covariates. Notes: Confidence intervals clustered by month of birth reported. Sample weighted to adjust for under sampling of less educated. All results adjust for month and year of birth, sex, and the ten principal components of population stratification.

**Supplementary Figure 7:**
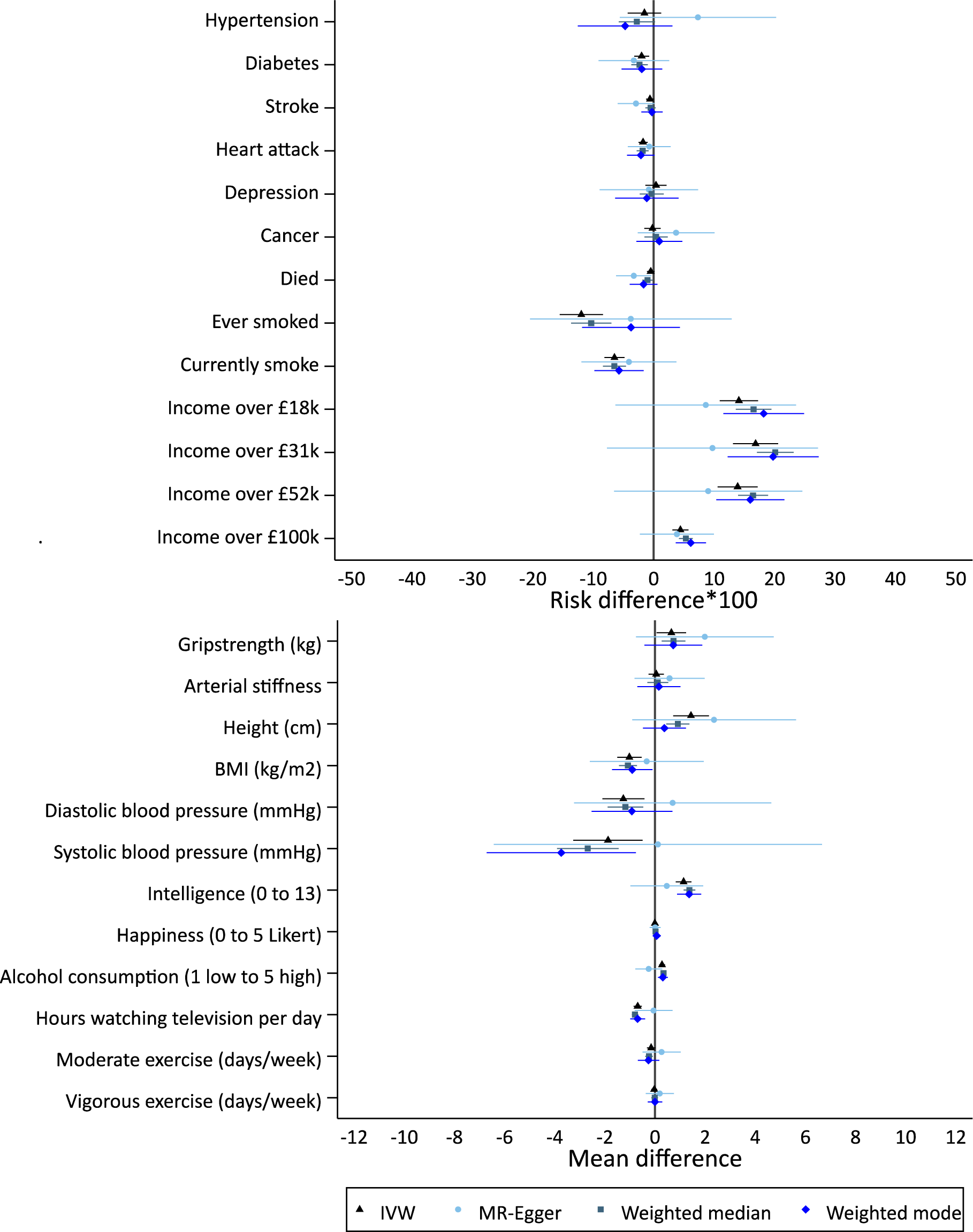
The effect of one additional year of schooling on morbidity, mortality and socioeconomic outcomes, estimated using the educational attainment GWAS genome-wide significant SNPs using inverse variance weighted, MR-Egger regression, weighted median and weighted modal estimators. Notes: Confidence intervals clustered by month of birth reported. Sample weighted to adjust for under sampling of less educated. All results adjust for month and year of birth, sex, and the ten principal components of population stratification. I^2^_gx_=0.21, this suggests that MR-Egger may be biased towards the null, as there is only modest variation in the SNP-educational attainment associations.

**Supplementary Table 1:** The estimated heterogeneity in the estimated effect of education across SNPs. Estimated using the I-squared statistic.(*43*)

| **Binary outcomes** | $I^2$ | 95% Confidence interval | |
| --- | --- | --- | --- |
|  |  | Lower | Upper |
| Hypertension | 0.610 | 0.499 | 0.697 |
| Diabetes | 0.470 | 0.303 | 0.596 |
| Stroke | 0.238 | 0.000 | 0.433 |
| Heart attack | 0.152 | 0.000 | 0.371 |
| Depressive episode | 0.451 | 0.277 | 0.583 |
| Cancer | 0.063 | 0.000 | 0.298 |
| Died | 0.179 | 0.000 | 0.391 |
| Ever smoked | 0.726 | 0.655 | 0.782 |
| Currently smoke | 0.462 | 0.292 | 0.591 |
| Income over £18k | 0.661 | 0.567 | 0.734 |
| Income over £31k | 0.771 | 0.715 | 0.816 |
| Income over £52k | 0.822 | 0.782 | 0.855 |
| Income over £100k | 0.735 | 0.667 | 0.789 |
| **Continuous outcomes** |  |  |  |
| Grip strength (kg)* | 0.795 | 0.746 | 0.834 |
| Arterial Stiffness* | 0.088 | 0.000 | 0.322 |
| Height (cm)* | 0.887 | 0.865 | 0.906 |
| BMI (kg/m2)* | 0.857 | 0.826 | 0.882 |
| Diastolic blood pressure (mmHg)* | 0.787 | 0.736 | 0.828 |
| Systolic blood pressure (mmHg)* | 0.744 | 0.679 | 0.796 |
| Intelligence (0 to 13)* | 0.829 | 0.791 | 0.861 |
| Happiness (0 to 5 Likert)* | 0.111 | 0.000 | 0.340 |
| Alcohol consumption (1 low, 5 high)* | 0.769 | 0.712 | 0.814 |
| Hours of television viewing per day* | 0.842 | 0.807 | 0.870 |
| Moderate exercise (days/week)* | 0.690 | 0.606 | 0.755 |
| Vigorous exercise (days/week)* | 0.558 | 0.427 | 0.660 |

**Author contributions**
NMD obtained funding for this study, analyzed and cleaned the data, interpreted results, wrote and revised the manuscript. MD interpreted the results, and wrote and revised the manuscript. GDS interpreted the results, wrote and revised the manuscript. FW interpreted the results, and wrote and revised the manuscript. GvdB interpreted the results, and wrote and revised the manuscript.

## Acknowledgements

The Medical Research Council (MRC) and the University of Bristol support the MRC Integrative Epidemiology Unit [MC_UU_12013/1, MC_UU_12013/9]. The Economics and Social Research Council (ESRC) support NMD via a Future Research Leaders grant [ES/N000757/1]. No funding body has influenced data collection, analysis or its interpretations. This publication is the work of the authors, who serve as the guarantors for the contents of this paper. This work was carried out using the computational facilities of the Advanced Computing Research Centre - http://www.bris.ac.uk/acrc/ and the Research Data Storage Facility of the University of Bristol - http://www.bris.ac.uk/acrc/storage/. This research was conducted using the UK Biobank Resource.

